# A draft genome assembly for the heterozygous wild tomato *Solanum habrochaites* highlights haplotypic structural variations of intracellular immune receptors

**DOI:** 10.1101/2022.01.21.477156

**Authors:** Kyungyong Seong, China Lunde Shaw, Eunyoung Seo, Meng Li, Ksenia V Krasileva, Brian Staskawicz

## Abstract

*Solanum habrochaites* LA1353 is a self-incompatible, highly heterozygous wild tomato that is a useful germplasm resource for the study of metabolism, reproduction and disease resistance. We generated a draft genome assembly with PacBio HiFi reads and genome annotations, which underscored the expansion of gene families associated with metabolite-production, self-incompatibility, DNA regulation and immunity. After manually curating intracellular nucleotide-binding leucine-rich repeat immune receptors (NLRs), we found that *S. habrochaites* LA1353 has a larger NLR inventory than other wild tomato species. A great number of heterozygous local copy number variations (CNVs) driven by haplotypic structural variations further expands the inventory, both enhancing NLR diversity and providing more opportunities for sequence evolution. The NLRs associated with local CNVs predominantly appear in the helper NLR (NRC)-related phylogenetic clades and are concentrated in a few physical NLR gene clusters. Synteny analysis points out that these genomic regions correspond to the known NLR clusters from which experimentally validated, functional NLRs, such as *Hero, Mi-1.2* and *Rpi-amr1*, have been identified. Producing and incorporating Resistance Gene Enrichment Sequencing (RenSeq) data across wild tomato species, we reveal that the regions with local CNVs might have been shaped nearly equally by recent NLR gains and losses, along with enhanced sequence diversification that diminishes one-to-one orthology between heterozygous alleles. Our analysis suggests that these genomic regions may have accelerated evolutionary dynamics for NLR diversity generation in *S. habrochaites* LA1353.

## Introduction

*Solanum habrochaites* LA1353 is a self-incompatible, heterozygous wild tomato that grows in a suboptimal environment at 2,650 meters above sea level in the Andes (Fig. 1; Fig. S1) (Bauchet and Causse, 2012; Broz, Randle, *et al*., 2017; Broz *et al*., 2021). Besides its unique morphological, metabolic and reproductive features (Bedinger *et al*., 2011; Fan *et al*., 2017; Kilambi *et al*., 2017; Tohge *et al*., 2020), genetic diversity of *S. habrochaites* associated with disease resistance has drawn researchers’ interest (Rick and Chetelat, 1995). Many resistance (R) genes identified from this species have been introgressed into the cultivated tomatoes (*Solanum lycopersicum*) (Chaudhary and Atamian, 2017). Novel disease resistance is continuously searched against diverse pathogens, including oomycete *Phytophthora infestans* (Li *et al*., 2011; Nowakowska *et al*., 2014; Copati *et al*., 2019), fungi *Botrytis cinerea* (Finkers *et al*., 2007) and bacteria *Pseudomonas syringae* (Bao *et al*., 2015; Thapa *et al*., 2015).

**Figure 1.**
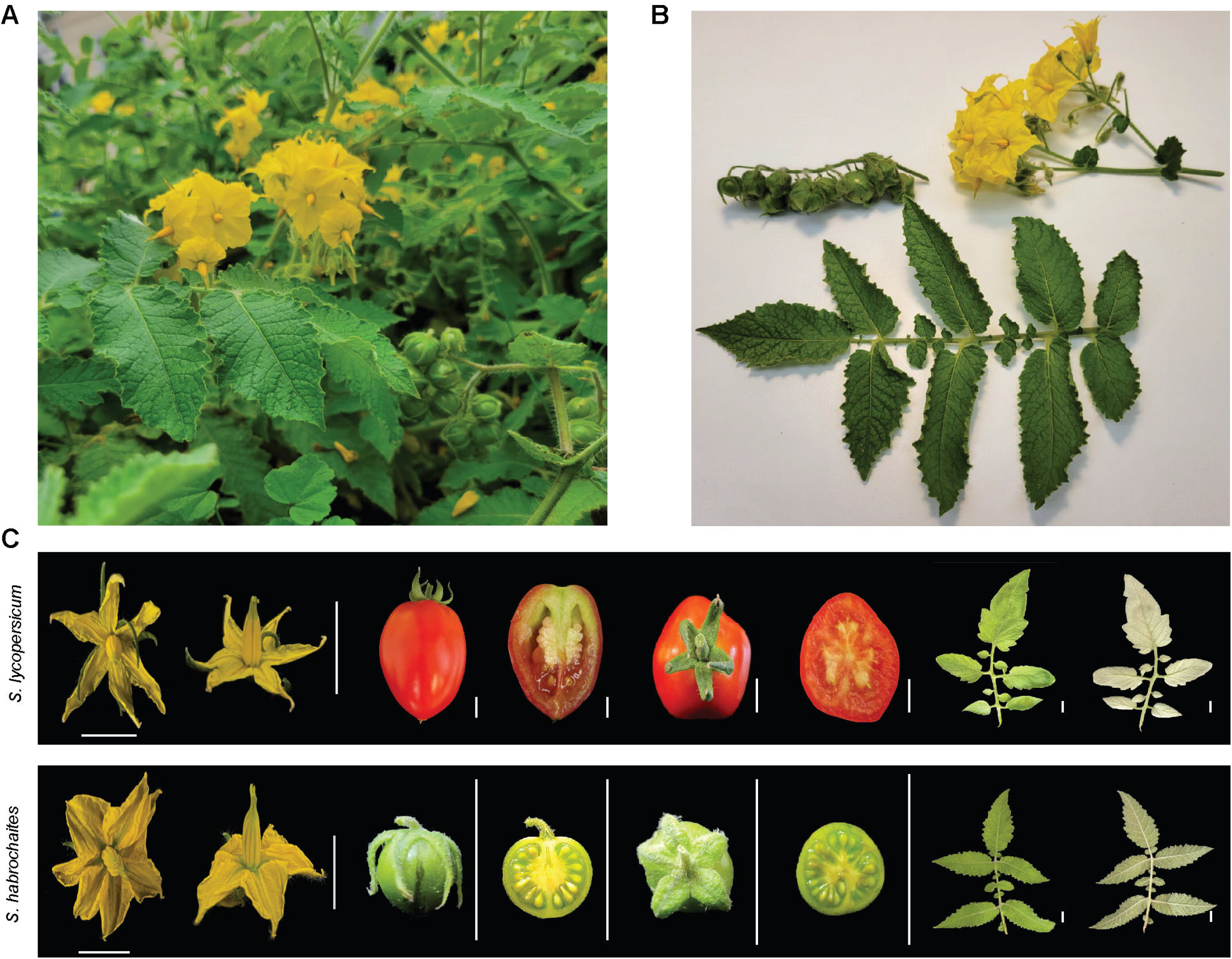
The morphological feature of *Solanum habrochaites* LA1353. **A**. S. *habrochaites* growing in the field at Berkeley, California, USA in November 2021. The plants were about three-month-old. **B**. Harvested S. *habrochaites* leaves, flowers and fruits. **C**. Flowers, fruits and leaves of *Solanum lycopersicum* Heinz and *S. habrochaites* LA1353. The white bars next to the organs indicate 1 cm. *S. habrochaites* fruits were about one month old.

Intracellular nucleotide-binding leucine-rich repeat immune receptors (NLRs) are essential components of the R gene-mediated immunity. Tomato and its wild relatives encode about 300 NLRs that have originated from diverse modes of duplications (Seo *et al*., 2016; Kim *et al*., 2017), and their multi-domain architecture (MDA) typically includes the central NB-ARC (NB) domain and C-terminal leucine rich-repeats (LRRs) in common, as well as N-terminal coiled-coil (CC), RPW8 domain or TIR domain. NLRs can be functionally classified as sensors that directly or indirectly perceive pathogen molecules (effectors) or helpers that execute a hypersensitive response in coordination with some sensors (Baggs *et al*., 2017). Sensors and helpers appear in distinct phylogenetic clades. In the CNL (CC-NB-LRRs) clades, the NRC clade members function as helpers with sensors in the NRC-dependent CNL superclade (Wu *et al*., 2017).

R gene enrichment sequencing (RenSeq) enables efficient study of NLRs through their selective amplification and sequencing from large genomes (Jupe *et al*., 2013; Witek *et al*., 2016; Stam *et al*., 2016; Seong *et al*., 2020). However, RenSeq outputs lack sufficient genomic contexts around NLRs. NLR clusters, as a result, tend to be fragmented into multiple contigs unless NLRs appear with close intergenic regions. Because of the missing context, it is infeasible to distinguish with RenSeq data of self-incompatible species heterozygous NLR alleles from recently duplicated NLRs in a haplotype. Thus, RenSeq data from highly heterozygous species does not align well with those from homozygous species that technically focus on haplotypic NLR variations. For this reason, our previous study relied only on the self-compatible, homozygous wild tomatoes and necessitated whole genome sequencing and assembly data for heterozygous wild tomatoes to investigate NLR evolution (Seong *et al*., 2020).

Although a genome assembly produced with Illumina short reads is available for *S. habrochaites*, it is highly fragmented and lacks annotations, making it difficult to utilize the assembly for understanding NLR evolution (The 100 Tomato Genome Sequencing Consortium *et al*., 2014). In this study, we generated a draft genome of *S habrochaites* LA1353 with PacBio High-Fidelity (HiFi) reads and annotations, which can be useful for diverse studies on this wild tomato. We report that copy number variations (CNVs) between haplotypes are concentrated in a few NLR gene clusters, which may be hotspots of NLR evolution in *S. habrochaites*.

## Materials and Methods

### RenSeq with single-molecule real-time sequencing on wild tomatoes

We obtained the seeds of *Solanum arcanum* LA2152, *Solanum huaylasense* LA1365, *Solanum peruvianum* LA0446, *Solanum corneliomulleri* LA1677, *Solanum chilense* LA1932, *Solanum pennellii* LA1272 and *S. habrochaites* LA1353 from the Tomato Genetics Resource Center (TGRC) at the University of California, Davis (Table S1). We used gDNA from young leaf tissues to perform RenSeq in combination with single-molecule real-time sequencing (SMRT), following the previously published protocol (Seong *et al*., 2020). The library was sequenced using the PacBio Sequel platform at the Vincent J. Coates Genomics Sequencing Laboratory at the University of California, Berkeley. We used ccs v6.0.0 to generate HiFi reads from the sequencing output (three full-passes and > 99% accuracy) (Table S1). 70 nucleotide sequences were trimmed from both ends of the HiFi reads with cutadapt v1.16 to remove barcodes and adapters, and the trimmed reads of each accession were assembled with HiCanu v2.2 (genomeSize=7M -pacbio-hifi) (Martin, 2011; Nurk *et al*., 2020).

### Whole genome sequencing, assembly and scaffolding

We conducted whole genome sequencing (WGS) for *S. habrochaites* LA1353 with 10X Genomics and PacBio sequencing. The 10X Genomics sequencing was performed with the gDNA collected from the same plant used for the RenSeq. After the DNA repair and BluePippin DNA size selection, we sequenced the gDNA as 150 bp linked-reads using the HiSeq X Ten platform at the University of California, Davis. The linked-reads were converted to interleaved reads with longranger v2.2.2 and assembled with Supernova v2.1.1 (https://www.10xgenomics.com).

We extracted gDNA from young leaf tissues of another plant and sequenced the gDNA using the Sequel II platform at the Vincent J. Coates Genomics Sequencing Laboratory. HiFi reads were generated with ccs v6.0.0 and assembled with hifiasm v0.12 (-l2) (Cheng *et al*., 2021). The purge_dups package was used to separate out remaining haplotigs from the primary assembly, which were then concatenated with the alternative contigs (Guan *et al*., 2020). We scaffolded the primary contigs with the 10X Genomics linked-reads, relying on ARCS v1.1.1 and LINKS v1.8.7 (Warren *et al*., 2015; Yeo *et al*., 2018). We removed contigs and scaffolds that represent chloroplast and mitochondrial DNAs based on the similarity to the known plasmid DNAs from *S. lycopersicum*.

### Structural and functional annotation and annotation quality assessment

To run BRAKER v2.1.5 and MAKER v3.01.03 for structural annotations (Cantarel *et al*., 2008; Brůna *et al*., 2021), we collected protein sequences of annotated Solanaceae species from Sol Genomics and paired-end transcriptome data for *S. habrochaites* LA0407, LA1223, LA1777, LA2098 and LA2119 from the Sequence Read Archive (SRA) (Table S2) (Mueller *et al*., 2005; Fernandez-Pozo *et al*., 2015; Pease *et al*., 2016; Broz, Guerrero, *et al*., 2017; Arnoux *et al*., 2021). We filtered the adapters and low-quality reads from the paired-end libraries with Trim Galore v0.6.4 (-q 20 --illumina --paired) and Cutadapt v2.4. The remaining reads were mapped to the genome with BWA-MEM v2.0pre1 (Vasimuddin *et al*., 2019). Additionally, we performed transcriptome assembly with Trinity v2.11.0 using the filtered libraries pooled for each accession (Grabherr *et al*., 2011). TransDecoder v5.5.0 was then used to select coding sequences (CDS) with complete open reading frames from super-transcripts (https://github.com/TransDecoder).

The *S. habrochaites* repeat library was generated with RepeatModeler v2.0.1 and used to soft-mask the assemblies with RepeatMasker v4.0.7 (Smit *et al*., 2013; Flynn *et al*., 2020). The mapped transcriptomic data and the collected protein sequences were used as evidence in BRAKER (--etpmode --softmasking), which relied on GeneMark v4.65_lic and AUGUSTUS v3.3.3 (Stanke *et al*., 2006; Brůna *et al*., 2020). We used MAKER to capture gene models supported by the assembled transcripts and protein evidence mainly relying on exonerate without ab initio predictors (est2genome=1; prot2genome=1) (Slater and Birney, 2005). The gene models with ≥ 98% bidirectional coverage against the top homologous hits in the protein annotation of *S. lycopersicum* (ITAG 4.1) were retained. A subset of these MAKER gene models with AED scores < 0.07 were used to train SNAP (Korf, 2004); randomly selected 2,000 gene models from the subset were used to train AUGUSTUS. We ran MAKER again with the external gene models from BRAKER (pred_model), as well as two ab initio predictors, SNAP and AUGUSTUS. The BRAKER and MAKER gene models with ≥ 98% bidirectional coverage against the top ITAG models were passed to the model_pred.

We sequentially selected final gene models from the exonerate-driven MAKER gene models and BRAKER gene models with ≥ 98% bidirectional coverage against the top ITAG annotations, MAKER gene models with AED scores ≤ 0.35 or InterPro matches, and BRAKER gene models supported by any hints. For this purpose, InterProscan v5.52-86.0 was used to search the sequences against Pfam v33.1, SUPERFAMILY v1.75 and Gene3D v4.3.0 (Wilson *et al*., 2009; Jones *et al*., 2014; Dawson *et al*., 2017; Mistry *et al*., 2021). Gene models composed only of transposable element (TE)-related domains were removed. The quality of the final annotation was assessed with BUSCO v5.2.2 and the Solanales datasets v10 (Kriventseva *et al*., 2019; Seppey *et al*., 2019).

### NLR manual curation

We identified putative NLR loci by translating the *S. habrochaites* genome assemblies in the six reading frames and searching for the NB-ARC domain with hmmsearch v3.3 (-E 1e-4 --domE 1e-4) (Eddy, 2011), as well as mapping RenSeq HiFi reads to the assemblies with BWA-MEM v2.0pre. We extracted the candidate loci with 10,000 flanking regions and manually annotated them as described previously (Seong *et al*., 2020). For other wild tomato species, we curated only a subset of contigs that contain NLRs belonging to the three phylogenetic clades of interest, G1, G8 and G14. These contigs were identified by searching the manually curated intact G1, G8 and G14 NLRs of *S. habrochaites* against the RenSeq contigs of wild tomatoes with exonerate v2.2.0 and selecting the best seven matches for each query (Slater and Birney, 2005).

### NLR classification

The NLRs were classified based on their MDA by the previously constructed pipeline (Seo *et al*., 2016). Gene models containing the NB-ARC domain were initially identified. Based on the N-terminal sequence homology to CC, RPW8 domain and TIR domain and the presence of C-terminal LRRs, the NLRs were classified to CNLs, RNLs, TNLs, NLs and Ns. We relied on NLR-parser to detect the major motifs on the NB-ARC domain and assigned an NLR as intact if its NB-ARC domain is equal to or larger than 160 amino acids and has three out of four major motifs present in order (Steuernagel *et al*., 2015). An NLR was classified as incomplete otherwise, and we did not analyze incomplete NLRs.

For phylogenetic classification, we collected experimentally validated NLRs in Solanaceae from RefPlantNLR (Kourelis *et al*., 2021). The NB-ARC domains of the intact NLRs from *S. habrochaites* LA1353, the reference NLRs and an outgroup CED-4 were aligned by MAFFT v7.313 (--globalpair --maxiterate 1000) (Katoh and Standley, 2013). The multiple sequence alignment was trimmed with TrimAl v1.4.rev22 (-gt 0.2) and used to infer a phylogenetic tree with RAxML v8.2.12 with 500 rapid bootstrapping (-p 12345 -x 12345 -m PROTGAMMAJTTF) (Capella-Gutierrez *et al*., 2009; Stamatakis, 2014). We then followed the previous study for phylogenetic grouping (Seong *et al*., 2020).

### Gene family analysis

We identified with OrthoFinder v2.5.4 (default) orthologous protein groups between *S. lycopersicum* Heinz (ITAG 4.1), *S. chilense* LA3111, *S. pimpinellifolium* LA2093, *S. pennellii* LA0716 and *S. habrochaites* LA1353 (Bolger *et al*., 2014; Hosmani *et al*., 2019; Stam *et al*., 2019; Emms and Kelly, 2019; X., Wang *et al*., 2020). We then used CAFE v5 to select orthologous groups that rapidly expanded in *S. habrochaites* in comparison to another tomato species (Mendes *et al*., 2021). The enrichment test was performed with the enricher function in clusterProfiler on PFAM domains of the expanded gene family members (Yu *et al*., 2012). Significant hits required p-value < 0.05 and q-value < 0.05. Any TE-related results were removed. For comparison of gene counts with PFAM domains of interest, we counted the genes only if the detected region of domain covers 60% or more of the full domain to prevent incompletely annotated gene fragments from inflating the counts.

## Results

### HiFi reads lead to good quality heterozygous genome assembly and annotation for *S. habrochaites* LA1353

We generated 175.1X of PacBio subreads and 56.6X of 10X Genomics sequencing data, given 2Gbp of the wild tomato genome size for both haplotypes (Table 1). From the subreads, 21.6 Gbp of HiFi reads were produced, providing 10.8X coverage for each haplotype in genome assembly. The final draft genome was 981.2 Mbp composed of 794 scaffolds with a N50 of 6.7 Mbp (Table 1). RepeatMasker detected 559.1 Mbp (57.0%) and 63.7 (6.5%) Mbp as retroelements and DNA transposons, respectively, indicating a high level of repeat content. *S. habrochaites* LA1353 is self-incompatible and heterozygous; consistently, GenomeScope estimated 1.31% of heterozygosity based on the 21-mer distribution of our Hifi reads (Fig. S2) (Marçais and Kingsford, 2011; Ranallo-Benavidez *et al*., 2020). The alternative assembly was 928.6 Mbp in size and consists of 6,306 contigs with a N50 of 524 kbp. Genome annotation captured 40,207 gene models from 268 scaffolds for primary assembly, and 35,412 gene models from 3,211 contigs for alternative assembly (Table 1). Each assembly showed 96.9% and 83.4% BUSCO completeness, collectively supporting high heterozygosity.

**Table 1.**
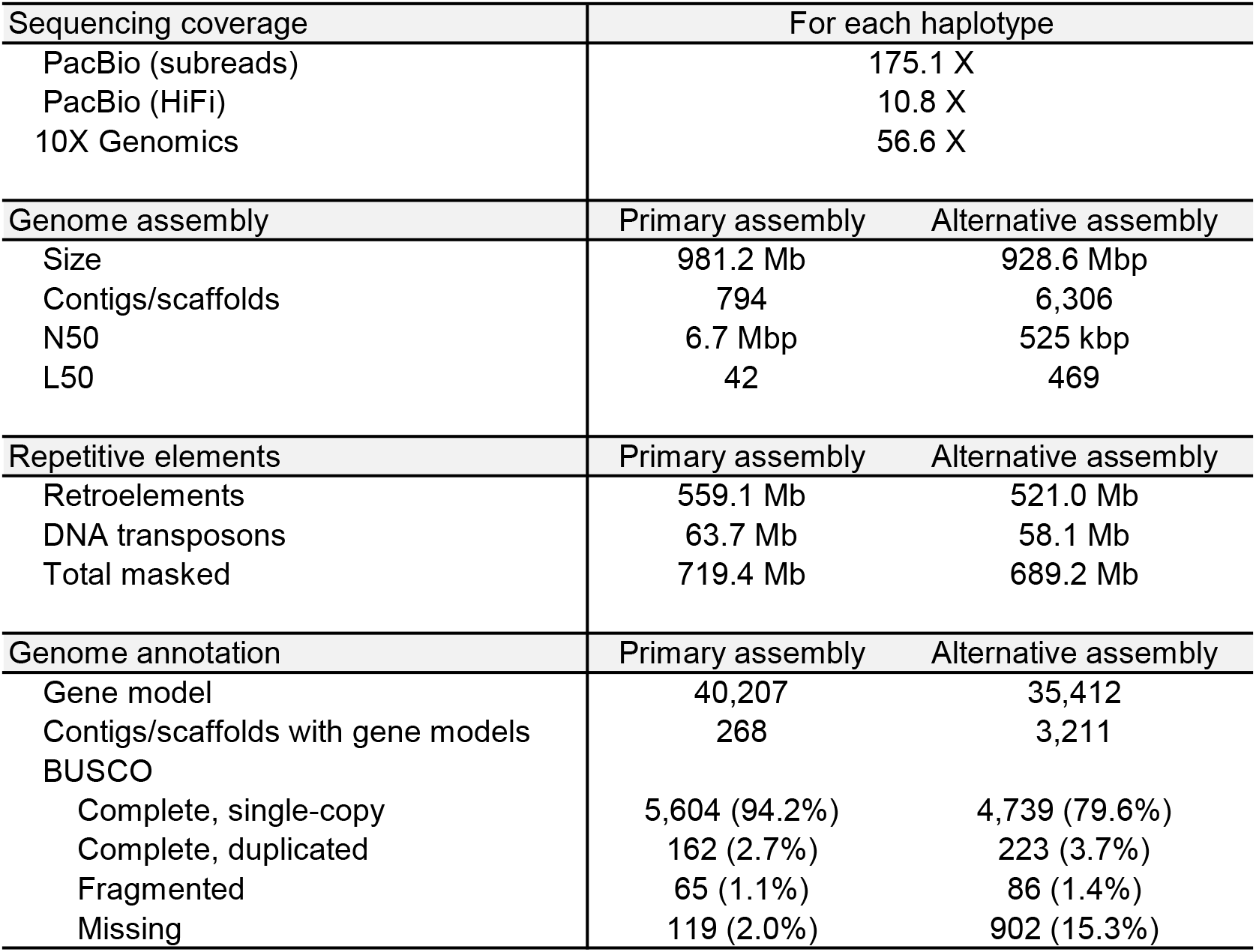
Genome sequencing, assembly and annotation statistics. Sequencing coverage was estimated based on the expected genome size of 2 Gbp for both haplotypes. Genome assembly statistics was calculated with QUAST (Gurevich *et al*., 2013). BUSCO was run on the protein annotation sets with the Solanales_odb10 database.

### The expanded gene families highlight the biological features of *S. habrochaites* LA1353

To examine whether our genome annotation supports the known biological features of *S. habrochaites*, we identified from the primary annotation set expanded orthologous groups and analyzed enriched PFAM domains (Fig. 2; Table S3). S. habrochaites is known as a rich source of disease resistance genes (Rick and Chetelat, 1995). Consistently, genes that contain domains found in NLRs, such as NB-ARC (PF00931) and Rx N-terminal domain (PF18052), and cell surface immune receptors, such as D-mannose binding lectin (PF01453) and S-locus glycoprotein domain (PF00954) were expanded. The number of self-incompatibility factors (PF05938) was the greatest in *S. habrochaites* LA1353, and DNA regulation and replication-associated domains were also enriched. *S. habrochaites* LA1353 develops long, dense trichomes and produces volatile metabolites that emanate a unique scent distinct from domesticated tomato, which together may function as protective barriers and repellents against pests (Bergau *et al*., 2015; F., Wang *et al*., 2020). Enzymes possibly involved in cell wall regulation and metabolite production were consistently expanded. Reflecting the adaptation of *S. habrochaites* LA1353 to a suboptimal environment at a high altitude of the Andes, Chlorophyll binding protein (PF00504) and genes associated with ATP production and usage appeared as significant hits.

**Figure 2.**
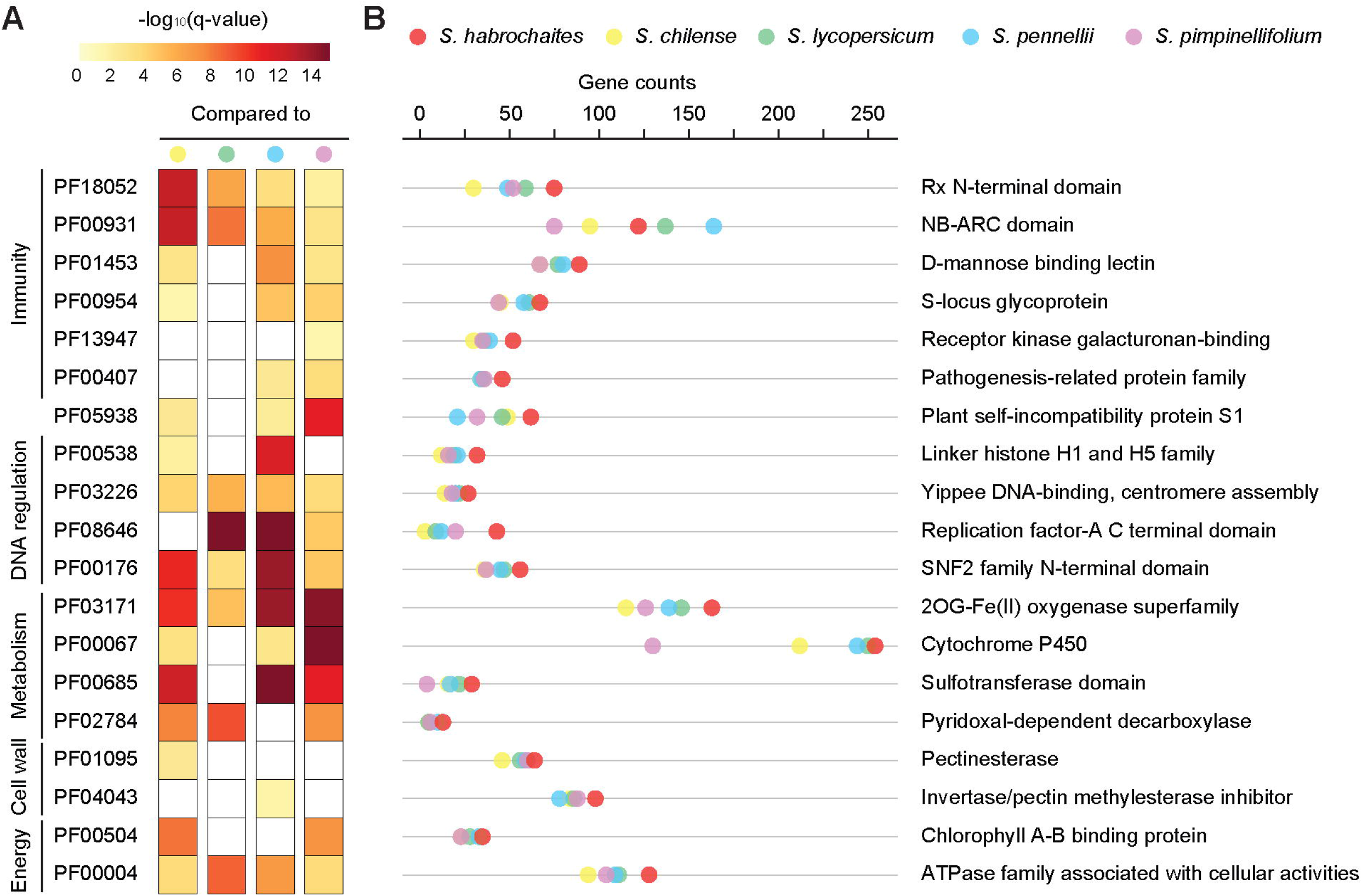
Expanded gene families in *S. habrochaites* LA1353. **A**. Enriched PFAM domains in rapidly expanded gene families identified in *S. habrochaites* LA1353 by CAFE. After orthologous groups were identified for all tomato species used for the analysis by OrthoFinder, the gene counts were compared in a pairwise manner between *S. habrochaites* LA1353 and another tomato species by CAFE to identify expanded orthologous groups in *S. habrochaites*. PFAM enrichment tests were performed for all rapidly expanded gene families using clusterProfiler. The cut-offs for p-value and q-value were 0.05, and the value of - log_10_(q-value) is capped at 15 for visualization. The empty boxes indicate no enrichment. **B**. The counts of the genes with the given PFAM entry. To prevent incompletely annotated gene fragments from inflating the gene counts, the genes were considered significant only if the predicted domain region is equal or larger than 60% of the full domain. Some of the PFAM domain names were simplified for visualization.

### The genome of *S. habrochaites* LA1353 displays high completeness for NLR loci assembly and annotation

Some NLRs proliferate through tandem duplication and with repetitive elements, forming complex NLR gene clusters (Kim *et al*., 2017; Barragan *et al*., 2019; Krasileva, 2019). Assembling such clusters and annotating these NLRs can be challenging for short reads and automatic annotation pipeline. For instance, we assembled 218 kbp of NLR gene clusters with HiFi reads in which we manually curated 18 NLRs and one NB-ARC fragment (Fig. 3A). In this region, automatic annotations fragmented, fused or only partially captured the NLR gene models. RepeatMasker classified some of the NLR loci as repeats (Bayer *et al*., 2018). Genome assembly we performed with 150 bp paired-end reads and Supernova assembler for the same accession failed to recover most NLR loci despite sufficient sequencing coverage.

**Figure 3.**
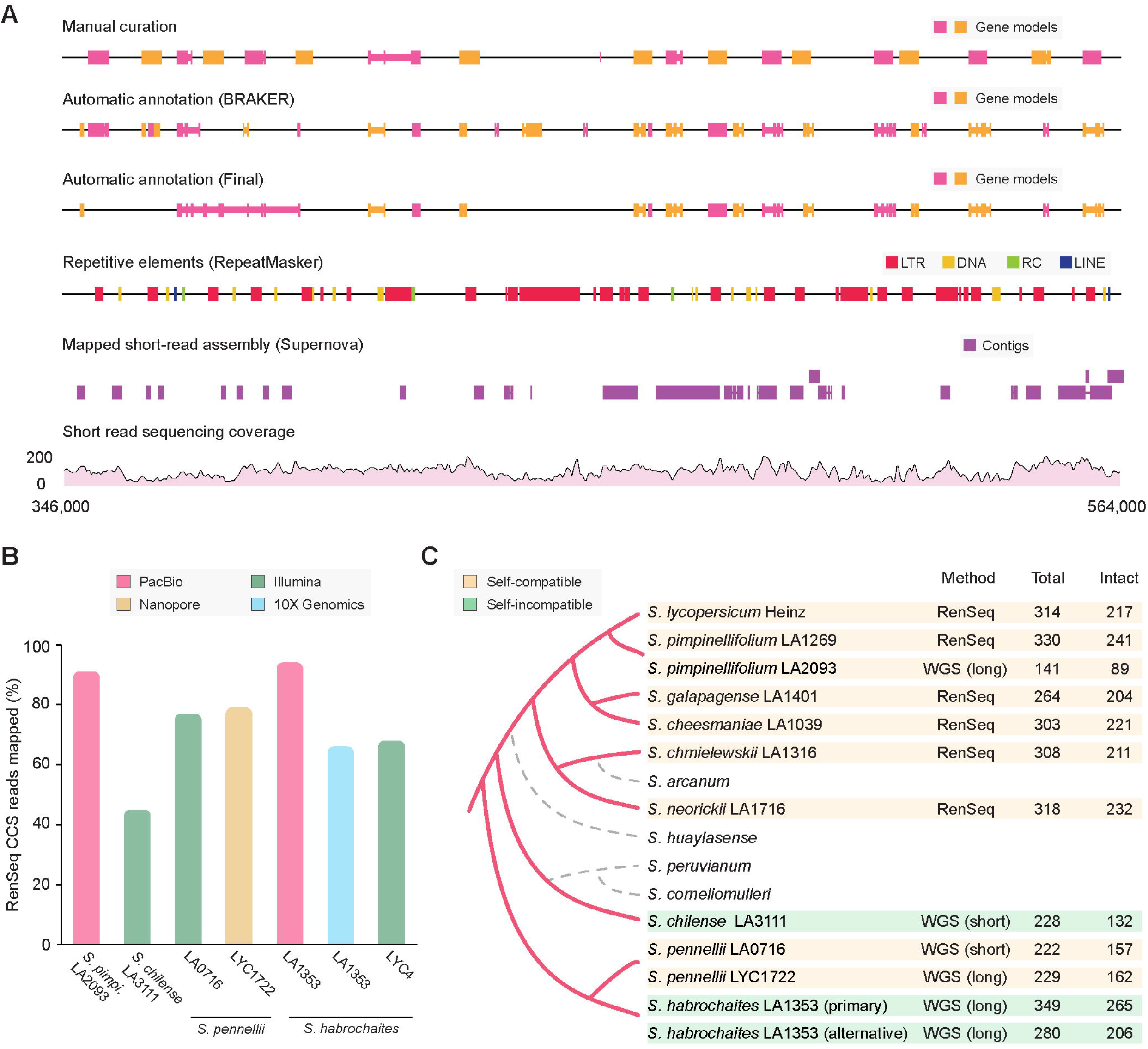
The completeness of NLR loci assembly and annotation of the *S. habrochaites* LA1353 draft genome. **A**. The 218 kb NLR gene cluster in scaffold 109 (346,000-564,000). 1st row: manually curated NLR gene models. Alternating colors are to indicate separate gene models. Exon and intron structures are distinguished by the height of the box, with larger boxes indicating exons. 2nd and 3rd rows: automatically annotated NLR gene models by BRAKER and in the final annotation set. 4th row: repetitive elements annotated by RepeatMasker. Only the known classes of transposons ≥ 300 nucleotides are indicated. 5th row: the contigs assembled with Supernova and 150 bp linked-reads from 10X Genomics sequencing mapped to the region. Supplementary alignments are not shown. 6th row: the sequencing coverage from 10X Genomics sequencing reads used for Supernova assembly. **B**. The estimated completeness of NLR loci assembly. The RenSeq data were obtained for *S. pimpinellifolium* LA0411, *S. chilense* LA1932, *S. pennellii* LA1272 and *S. habrochaites* LA1353. The RenSeq data typically display inconsistent coverage across NLR loci and also include non-NLR regions. Therefore, the HiFi reads were filtered to reduce redundancy, and only those that contain homologous sequences to the NB-ARC domain were selected and mapped. Of the total number of the reads, those mapped as primary alignments with the coverage 80% were counted. **C**. The hypothetical species tree reconstructed based on the previous study (Bedinger et al., 2011) and the number of putative NLRs and intact NLRs. The branch length is not on an evolutionary scale. The NLR annotations were obtained from either SMRT RenSeq or whole genome sequencing (WGS) and assembly with long reads, such as PacBio or Nanopore, and short reads, such as Illumina. The total gene model was identified primarily by searching for the gene models that contain the NB-ARC domain. Those that contain NB-ARC domain with three major motifs in order and ≥ 160 amino acids were counted as intact. Dashed lines indicate missing genome annotations or RenSeq data for the species.

We accessed the quality of NLR loci assembly for the selected wild tomato genomes by mapping non-redundant SMRT RenSeq data of wild tomatoes, which contained translated sequences homologous to the NB-ARC domain, to the respective genomes (Fig. 3B). Over 90% of the read mapping rate was reported for *S. habrochaites* LA1353 and *S. pimpinellifolium* LA2093 genomes produced with PacBio long reads. This *S. habrochaites* genome clearly displayed higher completeness than the one we generated with 10X Genomics sequencing reads and the one previously assembled with Illumina sequencing reads (The 100 Tomato Genome Sequencing Consortium *et al*., 2014). About 80% of the HiFi reads were mapped to the *S. pennellii* genomes. The heterozygous *S. chilense* genome assembled with Illumina reads only had 45%. Although accession-level NLR variations would affect the mapping statistics, this result suggested that our draft genome of *S. habrochaites* LA1353 displays high completeness for NLR loci assembly compared to other wild tomato genomes.

The structural annotation pipeline predicted 198 proteins with the NB-ARC domain from the primary assembly. However, we manually curated 349 gene models that contain the NB-ARC domain by correcting chimeric gene models and capturing unpredicted NLRs (Fig. 3A and 3C). Such improvement was significant given that *S. pimpinellifolium* LA2093 ended up with only 141 putative NLRs annotated from automatic gene prediction while the actual NLR count should likely be comparable to *S. pimpinellifolium* LA1269 from our previous RenSeq data (Fig. 3B) (Seong *et al*., 2020). As our gene models include putatively pseudogenized NLRs and NB-ARC fragments without proper MDAs, we selected 265 intact NLRs that have three or more major motifs over the NB-ARC domain equal to or longer than 160 amino acids. In the six tomato species studied with SMRT RenSeq, the intact NLR number varied from 204 to 241; the 265 intact NLR in *S. habrochaites* LA1353 is greater in number than those of other tomato species.

### Most heterozygous local copy number variations appear in NRC-clade-related phylogenetic groups

In heterozygous diploid genome assembly, one of the haplotypes builds a contiguous primary assembly together with collapsed, nearly homozygous regions; the other haplotype is separated into an alternative assembly (Fig. 4A) (Cheng *et al*., 2021). To investigate heterozygous NLR variations, we manually curated 206 intact NLRs from the alternative contigs (Fig. 3C). We then classified the entire intact NLRs into five categories based on the sequence similarity and phylogenetic relationship of the NLRs, as well as synteny between the primary and alternative assemblies (Fig. 4B; Table S4): (i) an NLR annotated only from a primary scaffold without any assembled alternative contigs is ‘common’ for both haplotypes; the two NLRs annotated from both primary and alternative assemblies with one-to-one orthology are (ii) ‘common’ if their CDS are identical or (iii) ‘genetically variable’ otherwise; NLRs annotated from either a primary scaffold or an alternative contig because the other has lost the NLR loci are characterized by (iv) ‘PAVs’ (presence/absence variations) or (v) ‘local CNVs’ if these NLR do not or do have a paralog within the 50 kb regions in 3’ or 5’ end, respectively.

**Figure 4.**
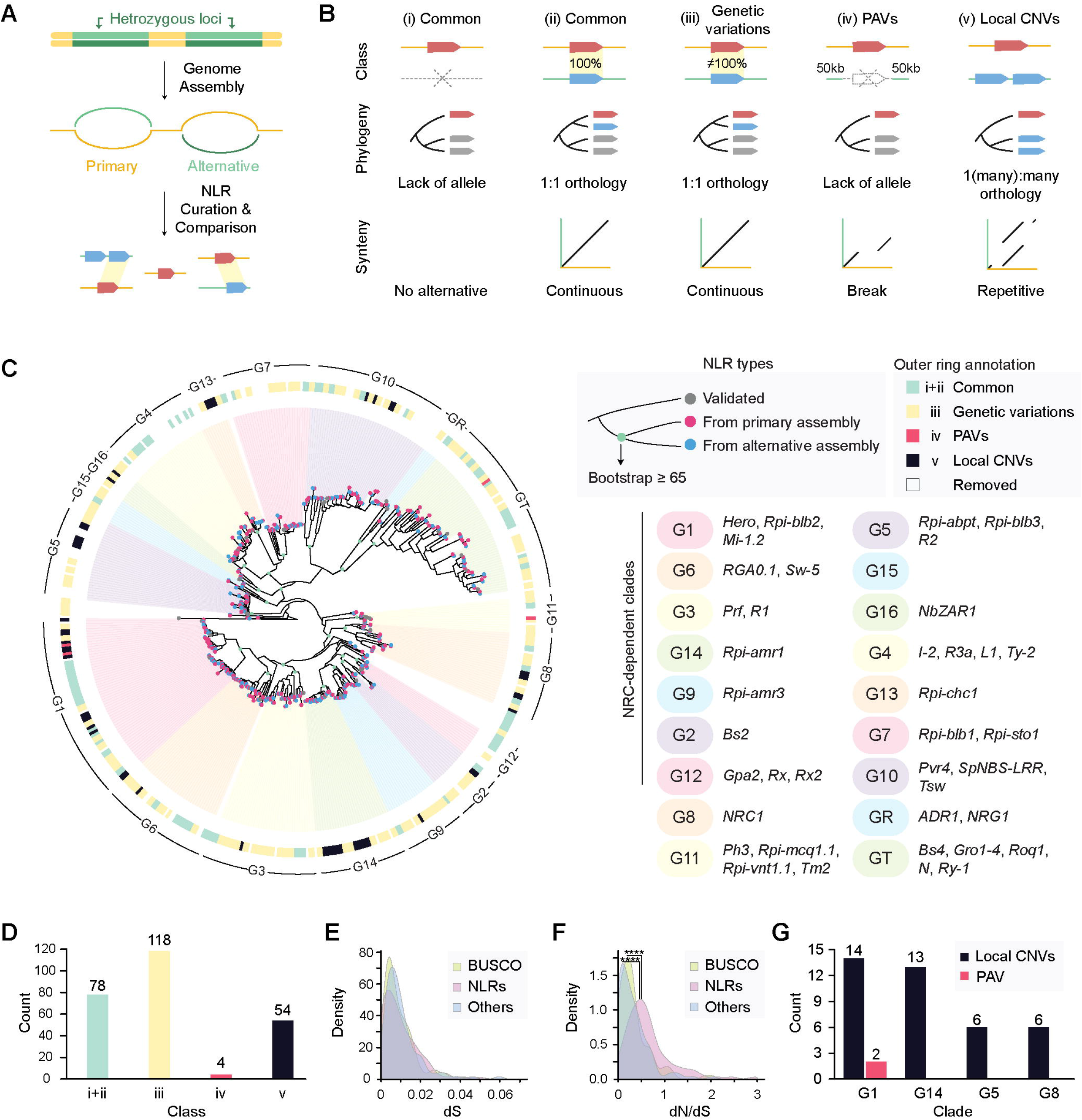
Heterozygous NLR variations in *S. habrochaites* LA1353. **A**. The schematics of heterozygous diploid genome assembly and NLR annotation. As haplotypes could not be fully resolved in the absence of parental DNAs, nearly or completely homozygous regions build contiguous primary assembly with one haplotype, while the other haplotype is separated to alternative assembly. **B**. The five categories of NLR variations. The provided phylogeny and synteny are representative but may not be the sole cases to explain each class. The phylogeny was created with GraPhlAn (Asnicar *et al*., 2015). **C.** A phylogenetic tree inferred for intact NLRs annotated from primary and alternative assemblies. The tree includes experimentally validated NLRs as references, and they are listed on the right. At the outer ring, the five classes given for each NLR are indicated with colors. **D**. The number of NLRs that belong to each class given in B. The colors correspond to the outer ring legend given in C. **E** and **F.** The distribution of synonymous substitution rates (dS) and nonsynonymous substitution rates (dN)/dS of three groups. 118 pairs of heterozygous alleles with one-to-one orthology were randomly selected from complete single-copy BUSCO genes or non-BUSCO genes that have at least one paralog in the genome (Others), following the size distribution of the genetically variable NLR pairs. According to the two-sided Kolmogorov-Smirnov test, the distribution of dS did not differ significantly, whereas the distribution of dN/dS was (P = 1E^−09^ for Others vs. NLRs and P = 2E^−10^ for Busco vs. NLRs). **G**. The number of PAVs and local CNVs found in the clades G1, G5, G8 and G14 that have the largest number of local CNVs.

During the classification, we removed any short or highly fragmented NLR gene models, because such gene structures are abnormal in comparison to closely related NLRs’ gene structures and may be indicative of pseudogenes. The majority of fragmented NLR gene models appeared in putative NRC-independent CNL clades, such as G4, G5, G7 and G10 (Fig. 4C and Table S4). Of the remaining gene models, 78 and 118 NLR pairs were common and genetically variable, respectively, and they appeared across all phylogenetic groups (Fig. 4D). To compare sequence evolution of the genetically variable NLR pairs, we randomly selected 118 pairs of heterozygous alleles from single-copy BUSCO genes or non-BUSCO genes that have at least one paralog in the genome. These heterozygous pairs had one-to-one orthology and followed a similar size distribution of the genetically variable NLRs. We then compared synonymous (dS) and non-synonymous (dN) substitution rates of these groups, respectively. All groups did not display noticeable differences in the dS distribution (Fig. 4E); however, increased dN significantly altered the dN/dS ratio of the NLRs, suggesting their unique haplotypic divergence (Fig. 4F). Besides, among heterozygous alleles were examples of frameshift mutations or transposon insertion only in one of the two alleles. Such events appeared across the phylogeny, which altogether possibly suggested distinct evolutionary dynamics of some heterozygous alleles.

Four and 54 NLRs were classified to have PAVs and local CNVs, respectively (Fig. 4D). This indicated that haplotype-specific gain or loss of NLRs not only is common but also preferentially appears in NLR gene clusters rather than isolated NLRs (Fig. 4B). Additionally, the local CNVs were phylogenetically biased (Fig. 4G). For instance, NRC-dependent CNL clades G1 and G14 had noticeably greater numbers of local CNVs than the other clades. For instance, despite the large clade size, no local CNVs were observed in the TNL group (GT) (Fig. 4C). Also, G3, G6 and G9 that are evolutionarily closely related to G1 and G14 only displayed single local CNV. Therefore, we conjectured that the frequent local CNVs in G1 and G14 are not simply attributed to the clade size but may be related to clade-specific NLR properties.

### A few known NLR gene clusters display most of the heterozygous local CNVs with complex evolution

The heterozygous local CNVs observed in clades G1, G5 and G14 are concentrated on a few NLR gene clusters, and these clusters correspond to the loci from which experimentally validated, functional NLRs were identified (Fig. 5A). For instance, a cluster in scaffold109 and two closely located clusters in scaffold47 together had all 14 CNVs found in G1. Based on synteny to the tomato reference genome, the NLR loci in these scaffolds corresponded to *Hero* and *Mi-1.2* clusters (Fig. 5A) (Vos *et al*., 1998; Ernst *et al*., 2002). Similarly, single clusters in scaffold102 and scaffold109 contained all 13 and 6 local CNVs of G14 and G5, respectively, and these loci corresponded to *Rpi-amrl* (G14) and *Rpi-blb3* (G5) gene clusters from wild potatoes (Lokossou *et al*., 2009; Witek *et al*., 2021).

**Figure 5.**
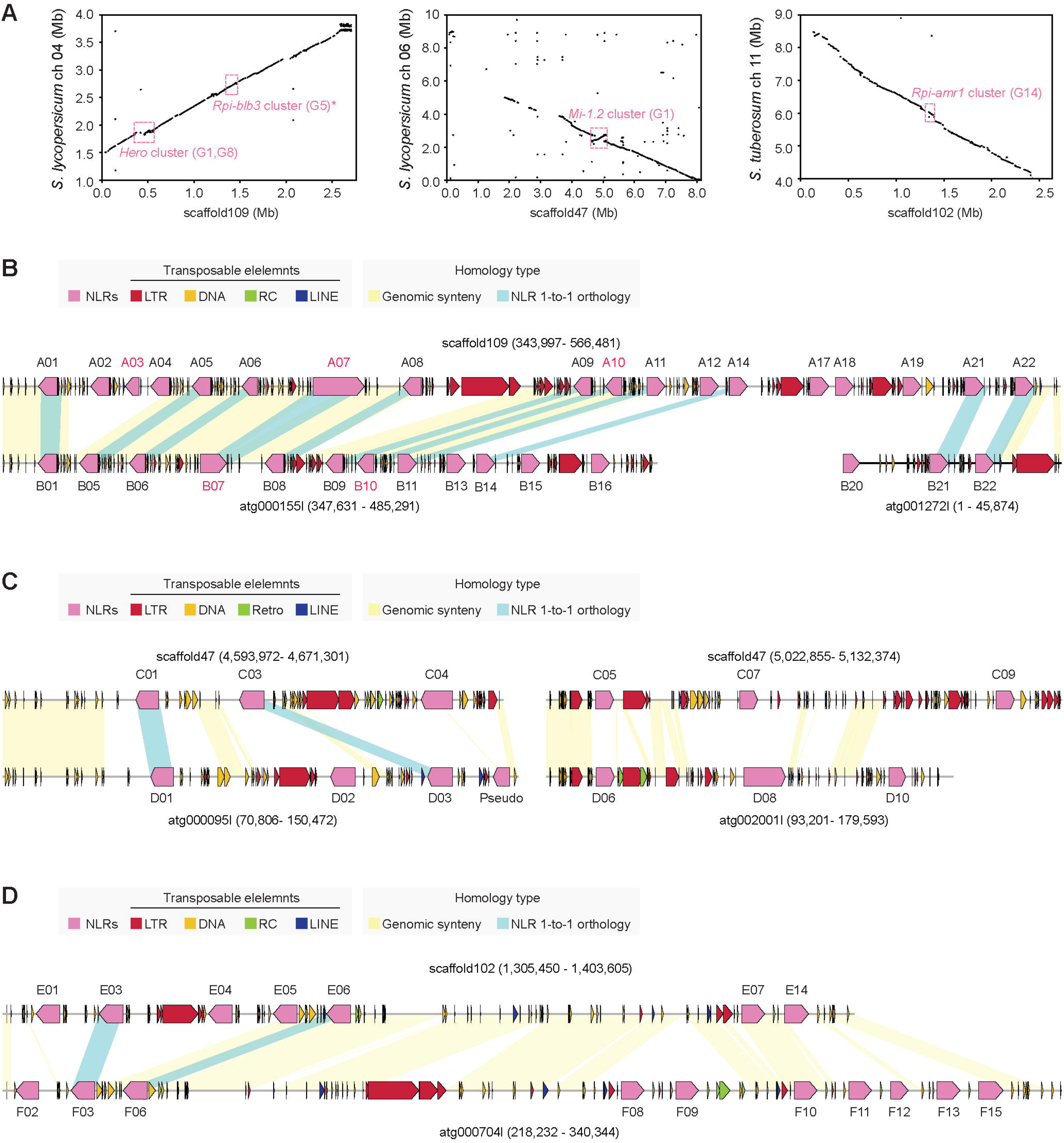
NLR gene clusters that display large numbers of heterozygous CNVs. **A**. Synteny between reference chromosomes of *Solanum lycopersicum* or *Solanum tuberosum* and primary scaffolds of *S. habrochaites* LA1353. Although the *Rpi-blb3* cluster is indicated in chromosome 4 of *S. lycopersicum*, this cluster was initially identified in potato. **B-D**. Schematics for NLR organization in the *Hero, Mi-1.2* and *Rpi-amr1* NLR clusters in *S. habrochaites* LA1353. The blocks indicate manually curated NLRs and transposable elements (TEs) annotated by RepeatMasker. The NLR blocks reflect gene lengths but not exon-intron structures; the TE blocks do not reflect the orientation. Genomic synteny was detected with progressiveMauve (Darling *et al*., 2010) and is highlighted in yellow. NLRs with one-to-one orthology were identified from phylogenetic trees (Fig. S4) and are highlighted in blue. The figures were generated with gggenomes (https://github.com/thackl/gggenomes). **B**. A17, A18 and A19 belong to the common class, as no alternative assembly that covers this region was found. Helper NLRs (G8) are indicated with red labels.

The synteny between homologous primary and alternative assemblies clearly indicated frequent homology breaks on the NLR loci (Fig. S3). Consistently, both the genetic organization and phylogenetic relationships of the NLRs with local CNVs hinted at complex evolution that has shaped these loci (Fig. 5B-D; Fig. S4; Table S5). Loss of one-to-one orthology appeared in all clusters. Some pairs, such as A12/B13, could be explained by accumulated mutation in a heterozygous allele and a recent duplication to B20 (Fig. 5B; Fig. S4A). In the *Mi-1.2* and *Rpi-amr1* clusters, only two heterozygous pairs have retained one-to-one orthology. C05/D06 and E01/F02 pairs, for instance, were phylogenetically distant although they occupy genomically proximal regions (Fig. 5C and 5D). The scenarios which could explain such findings are that the original orthologous allele was entirely swapped out by a paralogous NLR, or if attributed to sequence diversification, this would require more drastic processes like gene conversion through recombination. Phylogenetic distance is not necessarily congruent with physical distance (e.g. A12/A14 and C03/C04), suggesting that some paralogs did not result from duplication of the closest NLR (Fig. S4). Although differences exist in the TE profiles of primary and alternative contigs (Fig. 5B-D), we could not pinpoint particular TE-associated mechanisms that might have led to the cluster evolution.

### NLR gains and losses may nearly equally contribute to the cluster diversification

To examine how heterozygous NLR variations arose in the *Hero, Mi-1.2* and *Rpi-amr1* clusters, we incorporated interspecies NLR data to provide evolutionary contexts. The data include previously produced NLR annotations for six self-compatible tomato species (Fig. 2C) (Seong *et al*., 2020), as well as newly generated ones for heterozygous species with SMRT RenSeq (Table S1). As it is not possible to distinguish heterozygous NLR alleles from recently duplicated haplotypic paralogs, we focused on PAVs of orthologous NLRs in other tomato species. We then examined scenarios that can explain the architecture of the NLR clusters and phylogenetic relationships of NLRs with a minimum number of gene gains and losses (Fig. 6; Fig. S5).

**Figure 6.**
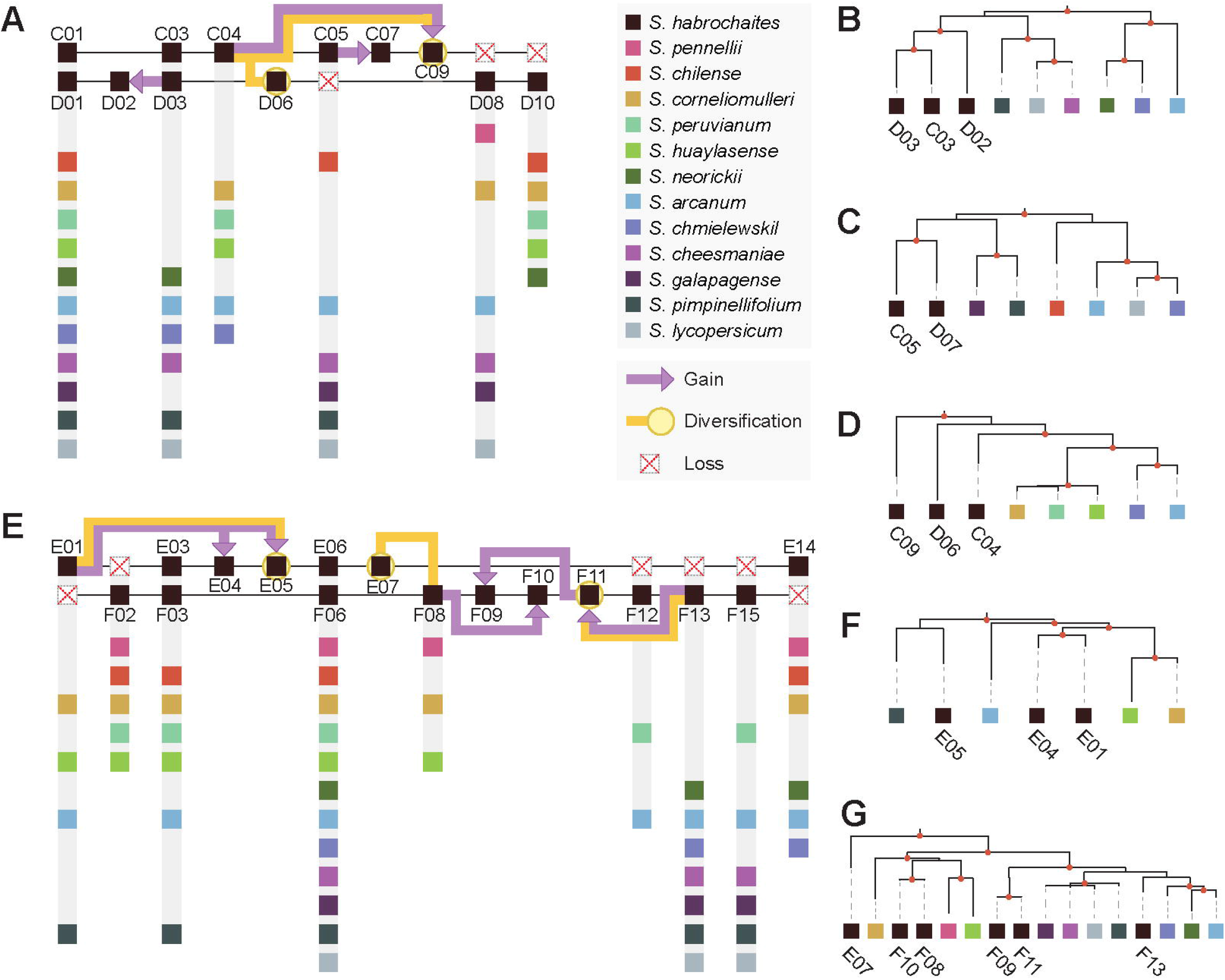
Interspecies NLR variations and suggested mechanism of heterozygous variations. **A** and **E**. Interspecies PAVs and evolution of *Mi-1.2* and *Rpi-amr1* clusters in *S. habrochaites* LA1353. NLRs in *Mi-1.2* and *Rpi-amr1* clusters are depicted in the hypothetical coordinates. The labels of NLRs are consistent with those in Figure 5. Orthologous genes were phylogenetically identified from other 12 tomato species, and their presence is indicated with colored boxes, regardless of the copy numbers. Putative evolutionary events—gain, loss and diversification—are mapped for NLRs from *S. habrochaites* LA1353 to explain haplotypic NLR variations and organization of NLRs. **B, C, D, F** and **G**. Simplified subsets of NLR gene trees. As RenSeq data for heterozygous species cannot be fully resolved, we focused on presence/absence variations. Therefore, one copy of NLR was chosen from the RenSeq data of each of the 12 tomato species, and the trees were pruned accordingly. The names are indicated only for NLRs from *S. habrochaites* LA1353. Red dots on the nodes indicate bootstrap ≥ 70.

The presence of orthologous genes in other tomato species suggests origin of the NLR prior to speciation. The absence of a heterozygous allele in this case can mean a haplotypic loss event (e.g., allele losses associated with D08 and F02 groups) (Fig. 6A and 6E; Fig. S5A). The presence of a paralogous copy with a monophyletic relationship with other *S. habrochaites* NLRs can point to recent haplotypic duplications (Fig. 6B, 6C, 6F and 6G). On the other hand, the emergence of phylogenetic singletons, such as C09 and E07, can be more plausibly attributed to duplication, followed by drastic sequence diversification, than independent multiple losses of orthologous alleles in other wild tomato species (Fig. 6D and 6G). In these scenarios, we explained the local CNVs with a minimum number of NLR loss and gain events. In the *Mi-1.2* cluster, 3 haplotypic NLR gains and losses were required to explain the heterozygous local CNVs. Similarly, 5 gains and 6 losses were mapped to the *Rpi-amr1* cluster, collectively suggesting that gains and losses of NLRs nearly equally contribute to diversifying the NLR clusters and shaping the local CNVs.

## Discussion

*S. habrochaites* is a useful germplasm source of disease resistance genes that could enhance resistance of domesticated tomatoes against diverse pathogens. RenSeq is a rapid and cost-effective approach to study disease resistance genes; however, its application to heterozygous species is not trivial and requires more careful consideration as RenSeq may not fully resolve complex NLR loci during assembly and annotation. In comparison to RenSeq data from homozygous species, the data from heterozygous ones generated two to three times the numbers of contigs and gene models, which contained redundant duplicates of one another. This was likely because RenSeq only captures a small subset of genomic regions, while resolving both heterozygous and haplotypic NLR loci requires more extensive genomic contexts. Therefore, elucidating NLR evolution in heterozygous species requires whole genome sequencing data.

Whole genome sequencing data alone, however, is not sufficient to study NLRs. Resolving complex NLR gene clusters depends on genome assembly with long, accurate reads and manual curation (Fig. 3A). In fact, the whole genome assembly we performed with short reads from 10X Genomics sequencing for the same accession failed to recover the majority of the *Hero* gene cluster. It is possible that masking NLR loci as repetitive elements and automatic annotation pipeline resulted in a low number of annotated NLRs in our genome and other long read assemblies (Fig. 3A and 3C). The quality of genome assembly and annotation would, therefore, lead to different conclusions about NLR evolution. For instance, previous studies have suggested that some wild tomatoes, such as *S. chilense* and *S. pennellii*, have lost a subset of NLRs (Stam *et al*., 2016; Stam *et al*., 2019). However, the relatively low mapping rates of our RenSeq data on the genomes assembled with short reads suggested that many NLR loci might have not been properly assembled (Fig. 3B). Therefore, we conclude that higher-quality genome assembly and annotation would be necessary to confirm the significant NLR loss events in certain lineages of wild tomatoes and convey a complete story of NLR evolution.

The *S. habrochaites* genome assembly presented in our study also captures drastic heterozygous NLR variations. Our study found that local CNVs appear more commonly than PAVs, indicating that structural variations are largely concentrated on NLR gene clusters, not isolated NLRs. Local CNVs predominantly appear in the NRC-related phylogenetic families. Two CNL sensor clades, G1 and G14, contain half of the NLRs associated with local CNVs (Fig. 4G). Although expected to have limited expansion (Wu *et al*., 2017), the conserved NRC helper clade (G8) had the third largest local CNVs. This could be due to co-evolution between sensors and helpers as hinted by the *Hero* gene cluster (Fig. 5B). All the local CNVs of G1 and G14 existed in *Hero, Mi-1.2* and *Rpi-amr1* (Fig. 5), which may have altered evolutionary dynamics with frequent duplications and rearrangement as suggested for *Arabidopsis* (Jiao and Schneeberger, 2020). Examining the PAVs of orthologous genes across 13 tomato species, we found that NLR gain and loss events may almost equally contribute to diversifying the gene clusters with possible accelerated sequence diversification for some loci (Fig. 6). Such processes diminish one-to-one orthology; the *Mi-1.2* and *Rpi-amr1* gene clusters, in particular, only have two pairs of NLRs that have maintained one-to-one orthology, suggesting each haplotype is possibly divergently evolving. The complexity of evolution suggested that events like unequal crossovers, gene conversions and strand slippage might be also actively involved in diversifying the NLR gene clusters (Barragan and Weigel, 2021). Overall, our data collectively suggests that highly heterozygous species often harbor two distinct NLR haplotypes; divergence of these haplotypes may provide more opportunities for heterozygous wild tomatoes to diversify resistance against pathogens.

The limitation of our data is that the haplotypes are not fully resolved. Future studies may sequence parental DNAs and generate haplotype-resolved genome assemblies with the trio-binning assembly. Controlled propagation of progeny and targeted sequencing of the *Hero, Mi-1.2* and *Rpi-amr1* clusters may also provide further insights into the evolution of complex NLR gene clusters. Additionally, not all *S. habrochaites* accessions are highly heterozygous. Unlike self-incompatible accessions like *S. habrochaites* LA1353 that populate southern Peru, the transition to self-compatibility is observed for the accessions found in central Peru and Ecuador (Broz, Randle, *et al*., 2017). Comparative genomics between self-compatible and self-incompatible *S. habrochaites* accessions could provide more contexts on how heterozygosity contributes to the NLR diversity generation.

## Data Availability

The *S. habrochaites* genome sequencing data, including PacBio raw reads, HiFi reads and linked-reads are available in the SRA in the National Center for Biotechnology Information (NCBI) under the BioProject accession PRJNA753882. Primary and alternative assemblies are deposited under PRJNA795504 and PRJNA795505. The HiFi reads from RenSeq can be accessed via PRJNA795506. All data, including genome assemblies, structural annotations and manually curated NLRs, can be accessed via Zenodo (10.5281/zenodo.5080564) or Sol Genomics Network (https://solgenomics.net).

## Acknowledgements

We thank the members at the genome center at the University of California Davis and the Vincent J. Coates Genomics Sequencing Lab at the University of California Berkeley for DNA extraction and sequencing. Seed was kindly provided by the Tomato Genetics Resource Center (TGRC) at the University of California, Davis. We appreciate the advice from Roger T. Chetelat at the TGRC on growing *S. habrochaites*. We thank Sarah Song for some photography and Erin Baggs and Pierre Joubert for reviewing the manuscript. This research relied on the Savio computational cluster resource provided by the Berkeley Research Computing program at the University of California, Berkeley. This work was supported by a grant from the Gordon and Betty Moore Foundation to the 2Blades Foundation (GBMF4725) and the Innovative Genomics Institute. Kyungyong Seong is supported by the Berkeley fellowship. Ksenia V Krasileva and China Lunde Shaw are supported by the Innovative Genomics Institute and NIH Director’s Award.

## Contribution

K.S. conceived and conducted the research and wrote the manuscript. M.L. performed DNA extraction and library preparation for RenSeq on the wild tomatoes. C.L.S. conducted pollination and field work. E.S. performed NLR classification. B.J.S. and K.V.K. supervised the research. All authors contributed to editing and reviewing this manuscript.

