## Supplemental Figures for "A draft genome assembly for the heterozygous wild tomato *Solanum habrochaites* highlights haplotypic structural variations of intracellular immune receptors"

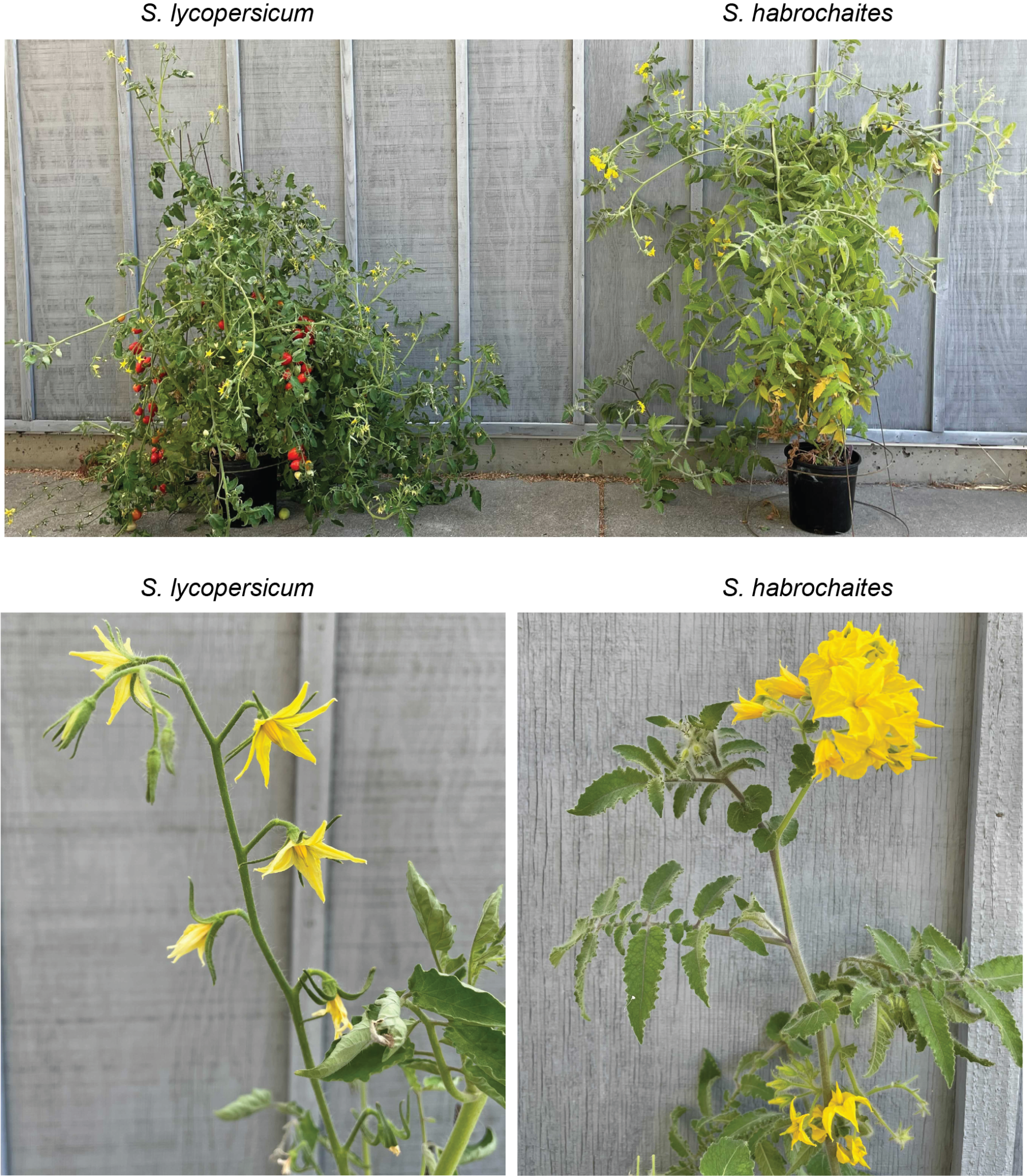

**Figure S1. Four-month-old *Solanum lycopersicum* and *Solanum habrochaites* SH1353 grown in a greenhouse from March to July.**

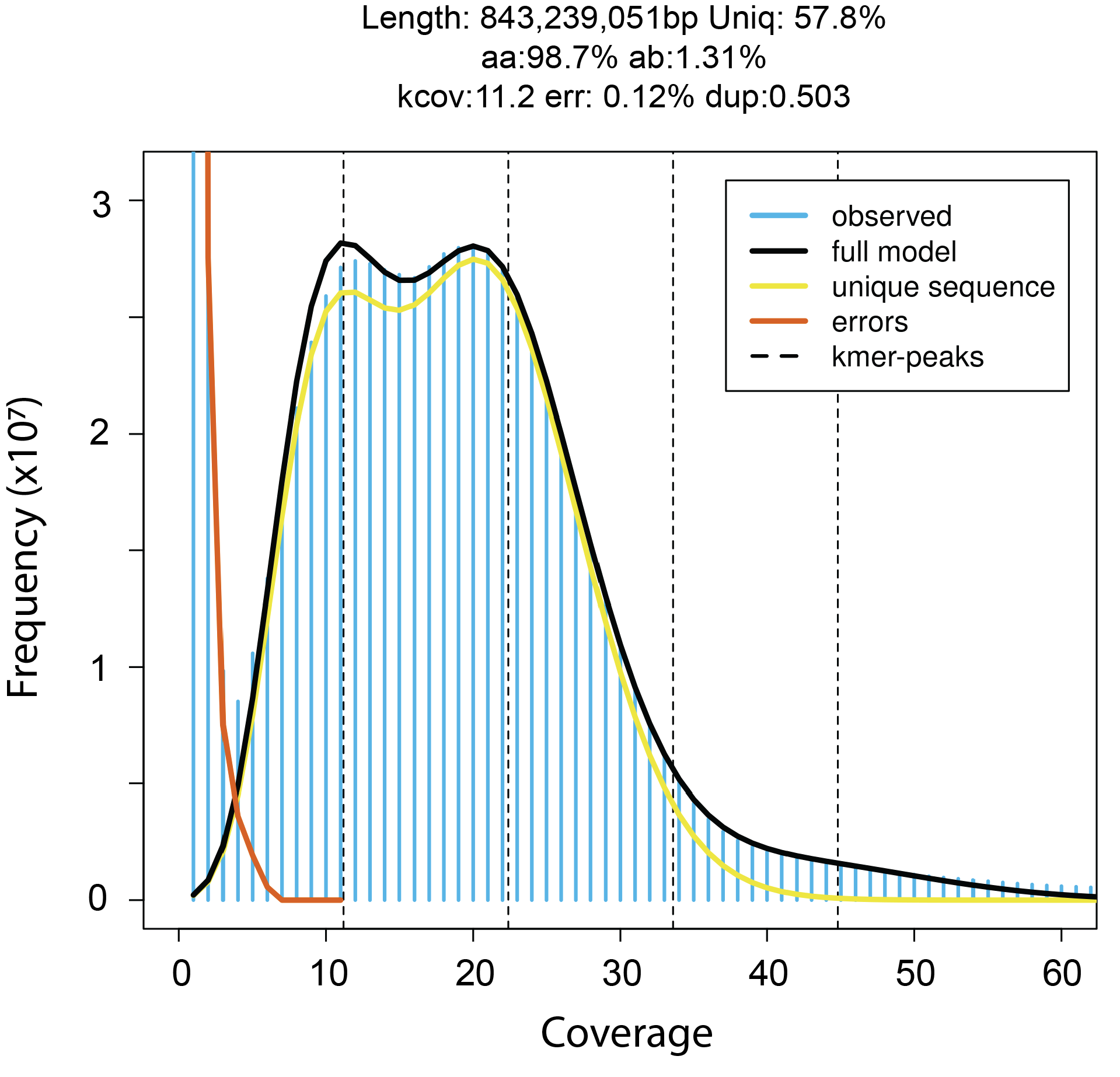

**Figure S2. The statistics of GenomeScope on the HiFi reads.**

GenomeScope was used to estimate heterozygosity and genome size with 21-mer.

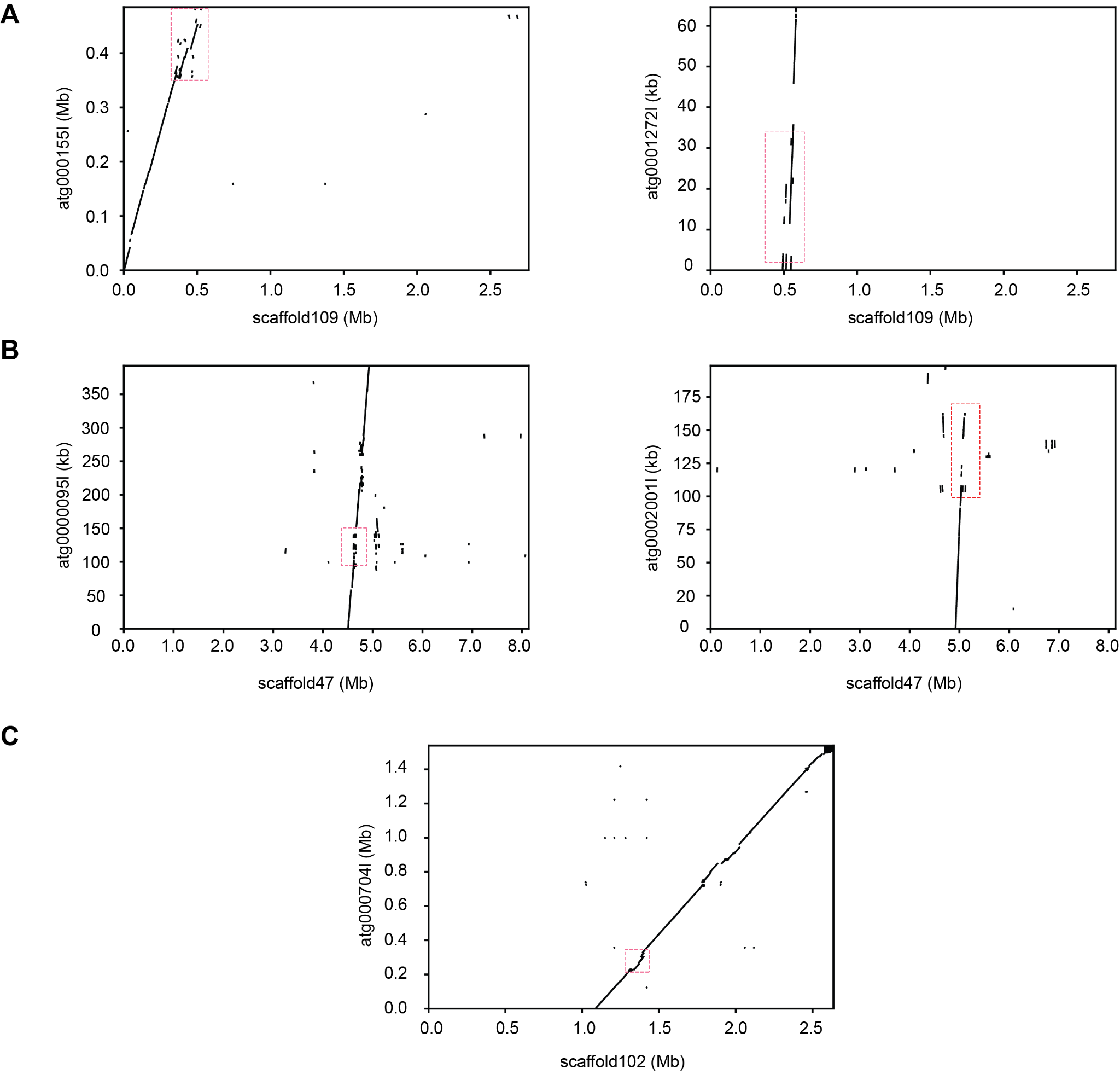

**Figure S3. Synteny between the homologous primary and alternative assemblies with local CNVs**

The homology detected with BLAST between primary scaffolds and their corresponding alternative contigs are visualized. The red boxes highlight the NLR loci with a great number of local CNVs. **A**. The *Hero* cluster. **B**. The *Mi-1.2* cluster. C. the *Rpi-amr1* cluster.

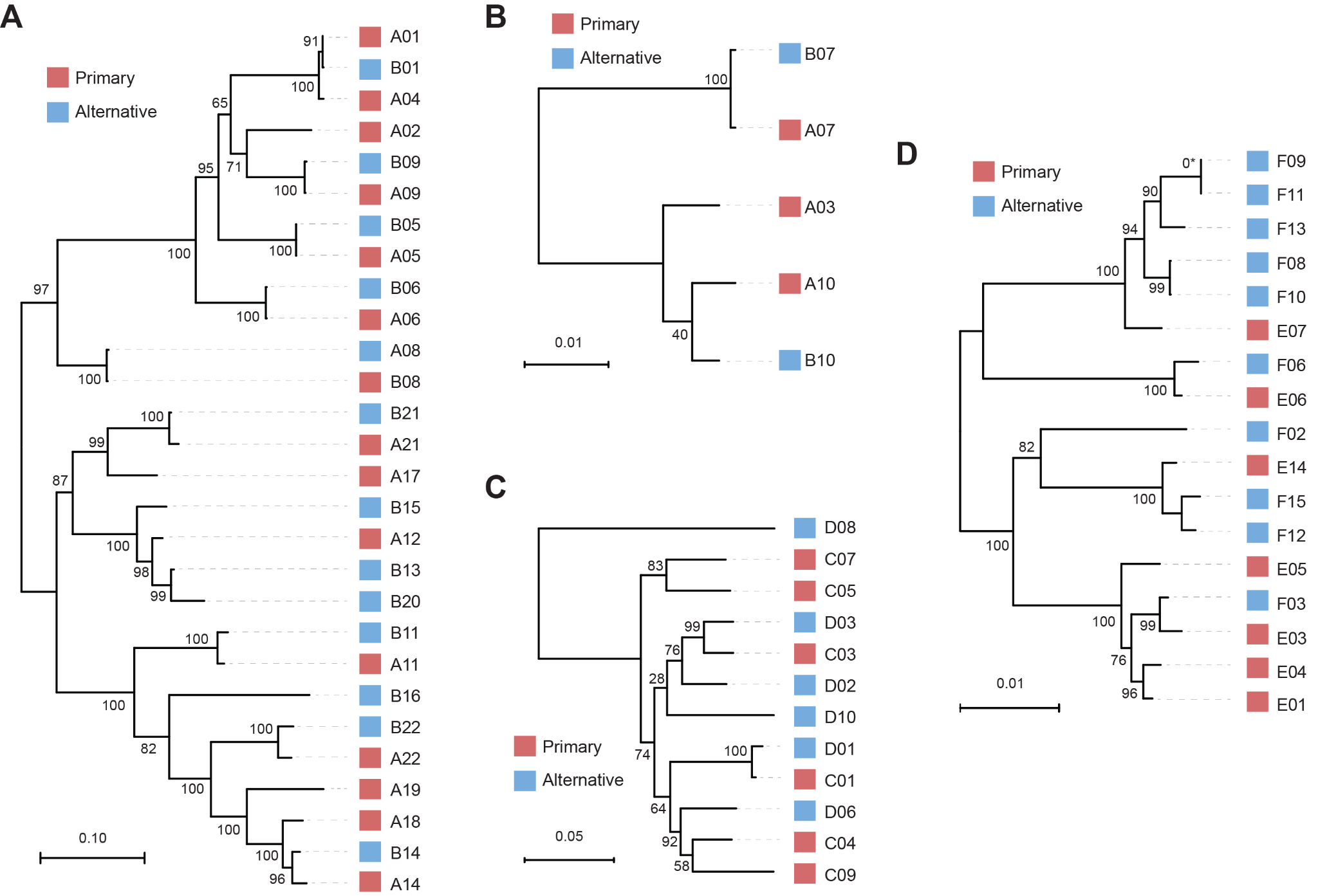

**Figure S4. The phylogenetic trees of NLRs in the selected NLR clusters**

The phylogenetic tree was inferred with the full-length NLRs in each cluster with FastTree and rooted at the midpoint. Red and blue boxes indicate NLRs that originated from primary scaffolds and alternative contigs, respectively. **A** and **B**. NLRs from the *Hero* cluster. Sensor NLR clade G1 (A) and helper NLR clade G8 (B). **C**. NLRs from the *Mi-1.2* cluster. **D**. NLRs from the *Rpi-amr1* cluster. The node of F09 and F11(*) is assigned with a bootstrap value of 0 as the two genes are identical.

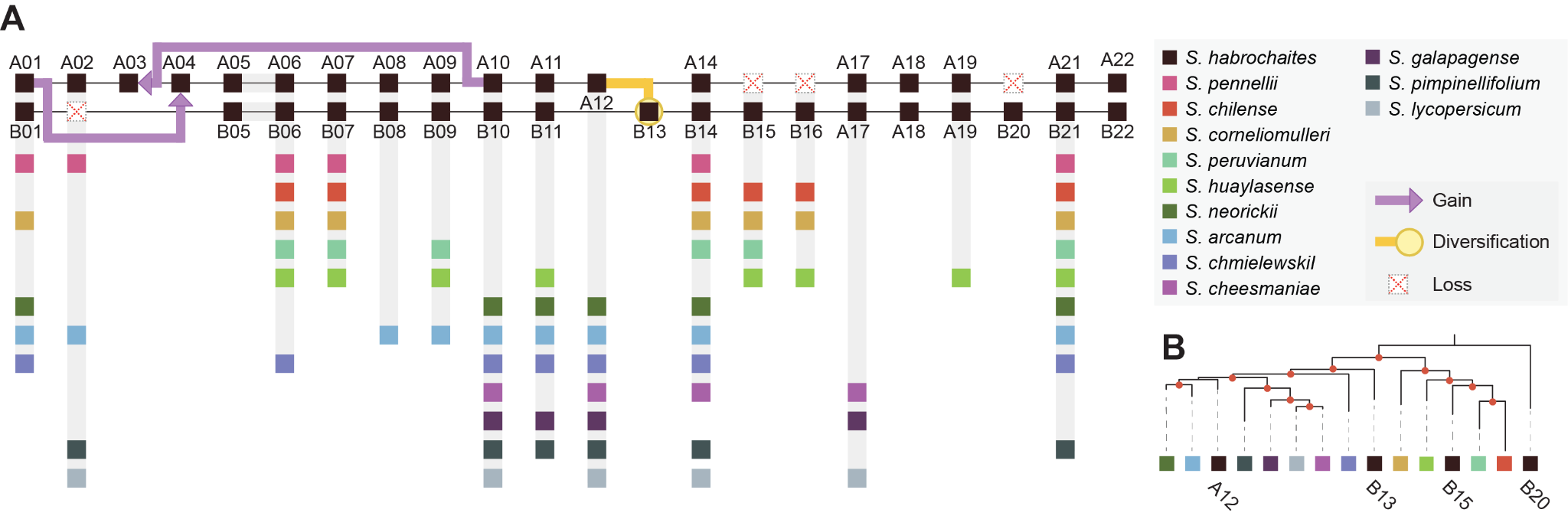

**Figure S5. Evolution of the *Hero* cluster**

**A.** Interspecies PAVs and evolution of the *Hero* cluster in *S. habrochaites* LA1353. NLRs in the *Hero* cluster are depicted in the hypothetical coordinates. Orthologous genes were phylogenetically identified from other 12 tomato species, and their presence is indicated with colored boxes, regardless of the copy numbers. Putative evolutionary events—gain, loss and diversification—are mapped for NLRs from *S. habrochaites* LA1353 to explain haplotypic NLR variations and organization of NLRs. **B.** Simplified subsets of NLR gene trees for those involved in sequence diversification. For simplicity, one copy of NLR was chosen from the RenSeq data of 12 tomato species, and the names are indicated only for NLRs from *S. habrochaites* LA1353. Red dots on the nodes indicate bootstrap ≥ 70.
